# Quality threshold evaluation of Sanger confirmation for results of whole exome sequencing in clinically diagnostic setting

**DOI:** 10.1101/2020.12.08.416792

**Authors:** Go Hun Seo, Hyeri Kim, Minjeong Kye, Jung-Young Park, Dong-gun Won, Jungsul lee

## Abstract

**Background:** With the ability to simultaneously sequence more than 5,000 disease-associated genes, next-generation sequencing (NGS) has replaced Sanger sequencing as the preferred method in the diagnostic field at the laboratory level. However, Sanger sequencing has been used routinely to confirm identified variants prior to reporting results. This validation process causes a turnaround time delay and cost increase. Thus, this study aimed to set a quality threshold that does not require Sanger confirmation by analyzing the characteristics of identified variants from whole exome sequencing (WES).

**Methods:** Our study analyzed data on a total of 694 disease-causing variants from 578 WES samples that had been diagnosed with suspected genetic disease. These samples were sequenced by Novaseq6000 and Exome Research Panel v2. All 694 variants (513 single-nucleotide variants (SNVs) and 181 indels) were validated by Sanger sequencing.

**Results:** A total of 693 variants included 512 SNVs and 181 indels from 578 patients and 367 genes. Five hundred seven heterozygous SNVs with at > 250 quality score and > 0.3 allele fraction were 100% confirmed by Sanger sequencing. Five heterozygous variants and one homozygous variant were not confirmed by Sanger sequencing, which showed 98.8% accuracy. There were 146 heterozygous variants and 35 homozygous variants among 181 indels, of which 11 heterozygous variants were not confirmed by Sanger sequencing (93.9% accuracy). Five non-confirmed variants with high quality were not identified on the ram .bam file.

**Conclusion:** Our results indicate that Sanger confirmation is not necessary for exome-derived SNVs with > 250 quality score and 0.3 > allele fraction set to an appropriate quality threshold. Indels or SNVs that do not meet the quality threshold should be reviewed by raw .bam file and Sanger confirmation should be performed to ensure accurate reporting.

## Introduction

Next-generation sequencing (NGS) is widely applied in the diagnosis of genetic diseases, as it allows the simultaneously sequencing of several millions of base pairs (1). When NGS is initially implemented for clinical diagnostic purposes, it is important to establish an optimization of the analytical specificity and sensitivity due to the complexity and limited experience in the clinical field. Clinicians or researchers have a sure degree of distress in accepting NGS results without re-confirming variants by Sanger method, which is viewed the gold standard for sequencing. Thus, Sanger sequencing has been routinely used to confirm identified variants prior to reporting the results (2).

However, the estimated cost of Sanger sequencing process for a single-nucleotide variant (SNV) is approximately US $240 and turnaround time is approximately one to one and a half weeks, including designing bi-directional PCR primers, running the analysis, analyzing chromatography, and interpreting the results (3). Conversely, as high quality NGS data are essential for clinical application, the accuracy of NGS results has been improved due to the establishment of quality assurance and quality programs in laboratories that perform NGS, as well as advances in technical processes of sequencing and bioinformatics analysis (2, 4–7). Several laboratories with different NGS sequencing platforms and sample sizes have determined quality thresholds for variants that do not need to be validate by Sanger sequencing to save costs and time (3, 8–12). SNVs with > 500 quality score did not require Sanger confirmation in 110 variants from 144 samples based on clinical exome sequencing (3). Another study reported that 7,601 variants from 5,109 samples that were analyzed using two NGS platforms and five targeted capture panels showed 100% validation rate in at least 35X depth coverage and more than 35% heterozygous ratio (8). A recently published study in 2019 was performed with 1,048 variants derived from exome sequencing and developed stringent criteria for variant calling (12). These efforts to reduce the range of Sanger confirmation have met expectations of improving turnaround time for reporting and reducing costs.

Since these quality thresholds could be unique according to their capture chemistry and analytical pipeline, a specified quality threshold for our analytical system should be set and validated. The purpose of this study was to set a quality threshold that does not require Sanger confirmation by analyzing the characteristics of the identified variants from whole exome sequencing (WES) that were performed in patients with suspected rare genetic disorders.

## Methods and materials

### Sample collection

Our study analyzed data on a total of 693 disease-causing variants. They were derived from WES of 578 individuals who had been diagnosed with suspected genetic diseases from May,2020 to October, 2020. Clinical manifestations of enrolled patients involved various systems. Patients’ samples were collected from individuals after obtaining the informed consent. Genomic DNA was extracted from the blood or buccal swab sample using the QIAamp blood kit (QIAGEN, GmbH, Germany) and AccuBuccal DNA Prep kit (AccuGene, Incheon, South Korea) protocol. The quality and quantity of DNA samples were measured using a NanoDrop spectrophotometer (Thermo Fisher Scientific, Waltham, MA, USA) and Qubit 4.0 fluorometer (Thermo Fisher Scientific, Waltham, MA, USA), respectively. DNA samples with an absorbance ratio A260/ A280 between 1.7 and 2.1 and a total amount greater than 1 μg were acceptable.

### Whole Exome sequencing

For the construction of standard exome capture libraries, we used the NEXTFLEX® Rapid DNA-Seq Kit 2.0 for Illumina paired-end sequencing library (Illumina, San Diego, CA, USA) and the xGen hybridization capture system with Exome Research Panel v2 (Integrated DNA Technologies, Coralville, IA, USA). The xGen Exome Research Panel v2 consists of 415,115 individually synthesized and quality-controlled xGen Lockdown Probes. The panel spans a 34 Mb target region (19,433 genes) of the human genome and covers 39 Mb of probe space, the genomic regions covered by probes.

Fragmentation of 100 ng of gDNA was performed using an E220 evolution focused-ultrasonicator (Covaris, Woburn, MA, USA). The fragmented DNA is repaired, an ‘A’ is ligated to the 3′ end, and adapters are then ligated to the fragments. Once ligation had been assessed, the adapter-ligated product was PCR amplified. The final purified product was then quantified and qualified using the TapeStation DNA screen tape (Agilent, Santa Clara, CA, USA). For exome capture, 500 ng of DNA library was mixed with hybridization buffers, blocking mixes, and xGen Exome Research Panel v2 probe, according to the standard xGen hybridization capture system protocol. Hybridization to the capture baits was conducted at 65°C using for 16 h on a PCR machine. The captured DNA was amplified. The final purified product was quantified using Qubit 4.0 and qualified using the TapeStation DNA screen tape (Agilent, Santa Clara, CA, USA). Sequencing was conducted on NovaSeq6000 using 150-bp paired-end conditions according to the manufacturer’s standard workflow (Illumina, San Diego, CA, USA).

### Bioinformatic analysis and variant selection

The quality of FASTQ files obtained by sequencing with the Illumina Novaseq 6000 was assessed using FASTQC (http://www.bioinformatics.babraham.ac.uk/projects/fastqc/). Subsequently, the base and sequence adapters with low base quality were removed using Trimmomatic. Pre-processed FASTQ files were aligned to the reference sequence (original GRCh37 from NCBI, February 2009) using BWA-MEM (v.0.7.17). Aligned BAM files were sorted and extracted using the statistical metric by samtools (v.1.9). Duplication was marked by Picard (v.2.20.8) (http://broadinstitute.github.io/picard/). SNVs and indel variants were called by HaplotypeCaller of GATK (v.3.8). Finally, variant call formats were generated. EVIDENCE developed by 3billion, Inc. was used to prioritize the variant according to the ACMG guideline and symptom similarity score as previously described (13). Relevant candidate variants including pathogenic variants, likely pathogenic variants, and variants of unknown significance based on EVIDENCE, were manually reviewed and selected for validation using Sanger sequencing. In total, 513 SNVs and 181 indels were selected for Sanger confirmation.

### Sanger Sequencing

The tested variant was amplified by PCR using primers designed with primer3 cgi v.3.0, Whitehead Institute (http://bioinfo.ut.ee/primer3-0.4.0/) and a reference sequence (NCBI GenBank). DNA was amplified using the PCR Master Mix Kit (ThermoFisher Scientific, Waltham, MA, USA) with the following conditions: 95°C for 5 min; followed by a program of 95 °C for 30 s, 60 °C for 30 s, and 72 °C for 30 s for 35 cycles; and ending with a 5 min extension at 72 °C. Amplicons were purified using ExoSAP-IT and bidirectionally sequenced using Big Dye Terminator version 3.1 on a SeqStudio Genetic Analyzer (Applied Biosystems, Foster City, CA, USA).

## Results

### The characteristics of identified variants from WES

The mean depth coverage was about 125 ± 24.5 in a total of 693 variants including 512 SNVs and 181 indels from 578 patients and 367 genes. Detailed information about the identified variants is shown in Supplementary Table 1. Allele frequency of 694 variants in the general population was <0.001 (https://gnomad.broadinstitute.org/). These variants and genes were distributed in different chromosomes of the human genome.

### Assessment of Sanger validation of identified variants from WES

Among the 512 SNVs, there were 433 heterozygous variants and 79 homozygous variants. Five heterozygous variants and one homozygous variant of them were not confirmed by Sanger sequencing, which translates into 98.8% accuracy. In the relationship between the quality score and the allele fraction (altered allele/reference allele +altered allele) of 512 variants, 417 heterozygous SNVs with > 250 quality score and > 0.3 allele fraction were confirmed by Sanger sequencing (Figure 1). Seventy-eight homozygous SNVs with a > 250 quality score and > 0.95 allele fraction were confirmed. The average quality score of six non-confirmed variants was 66.77 (Table 1)

**Figure 1.**
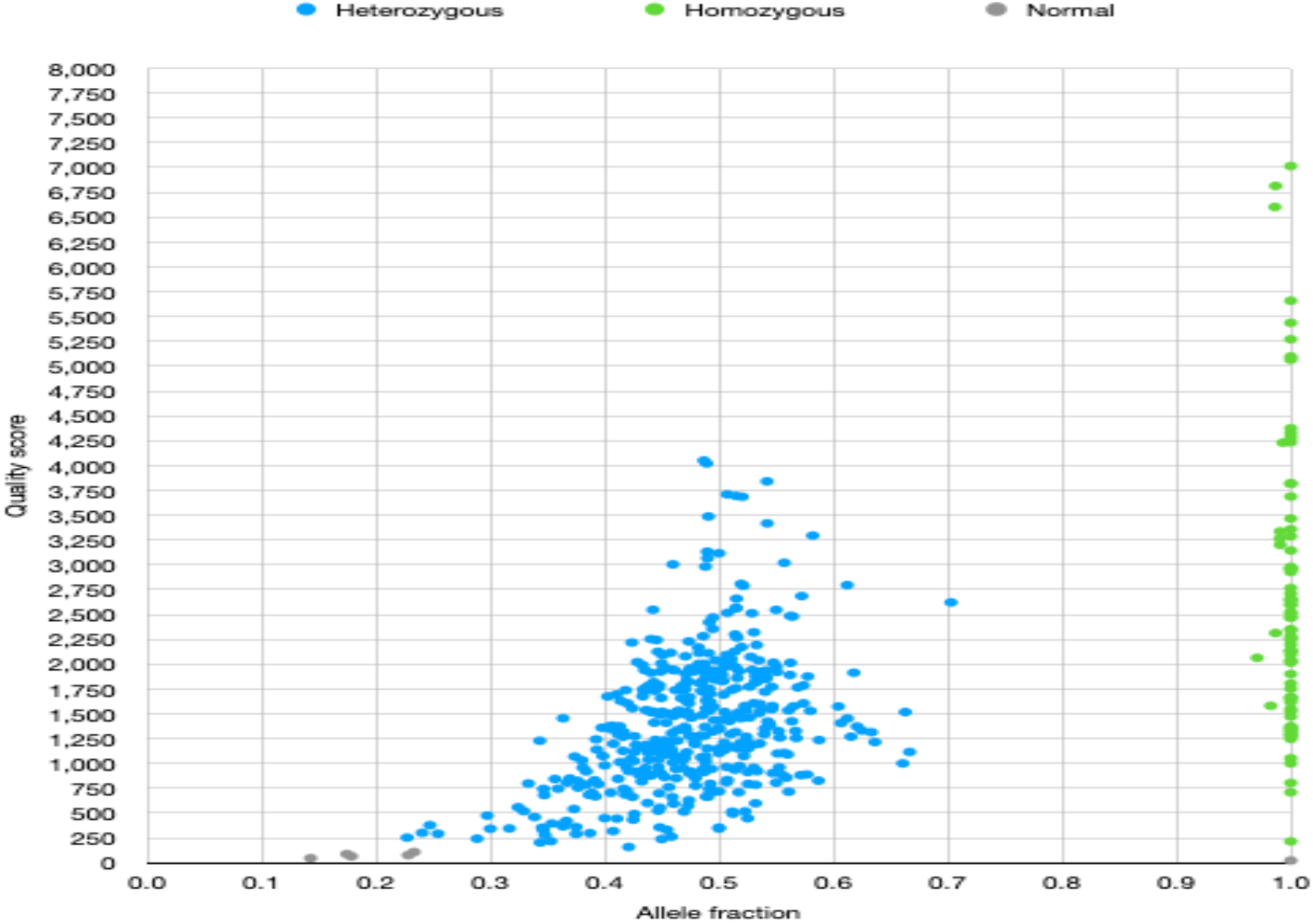
Distribution of SNVs according to the quality score and allele fraction

**Table 1.**
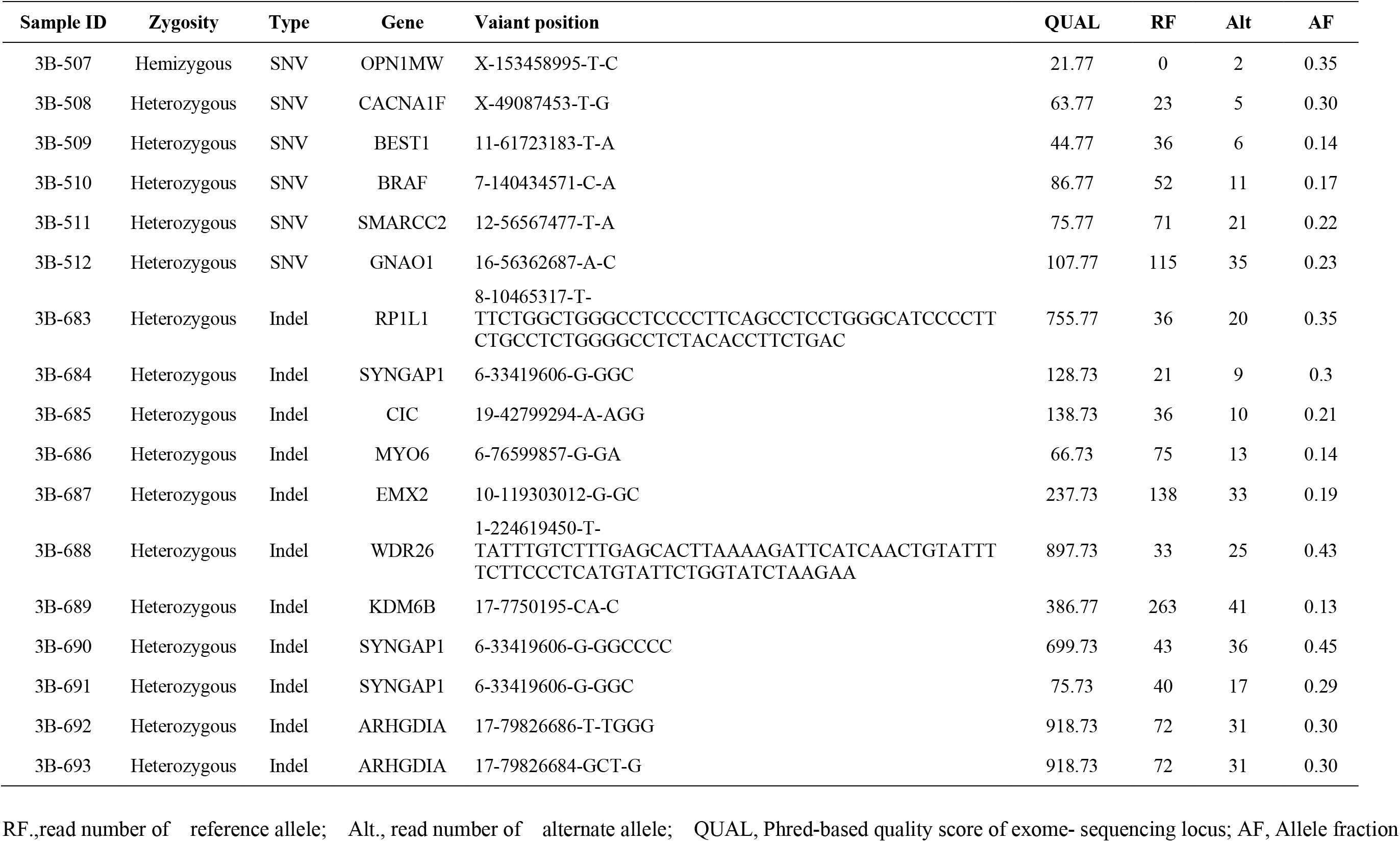
Detailed information on SNVs and indels not confirmed by Sanger sequencing

There were 146 heterozygous variants and 35 homozygous variants among 181 indels, of which 11 heterozygous variants were not confirmed by Sanger sequencing (93.9% accuracy). Of the146 heterozygous indels, 128 indels (87.6%) were confirmed by Sanger sequencing with a quality score of 250 or higher and 0.3 or higher for allele fraction (Figure 2). All homozygous indels were confirmed at a quality score of 250 or higher and 0.95 or higher. Five non-confirmed variants with >500 quality scores and >0.3 allele fractions were not identified on the ram .bam file (Table 1).

**Figure 2.**
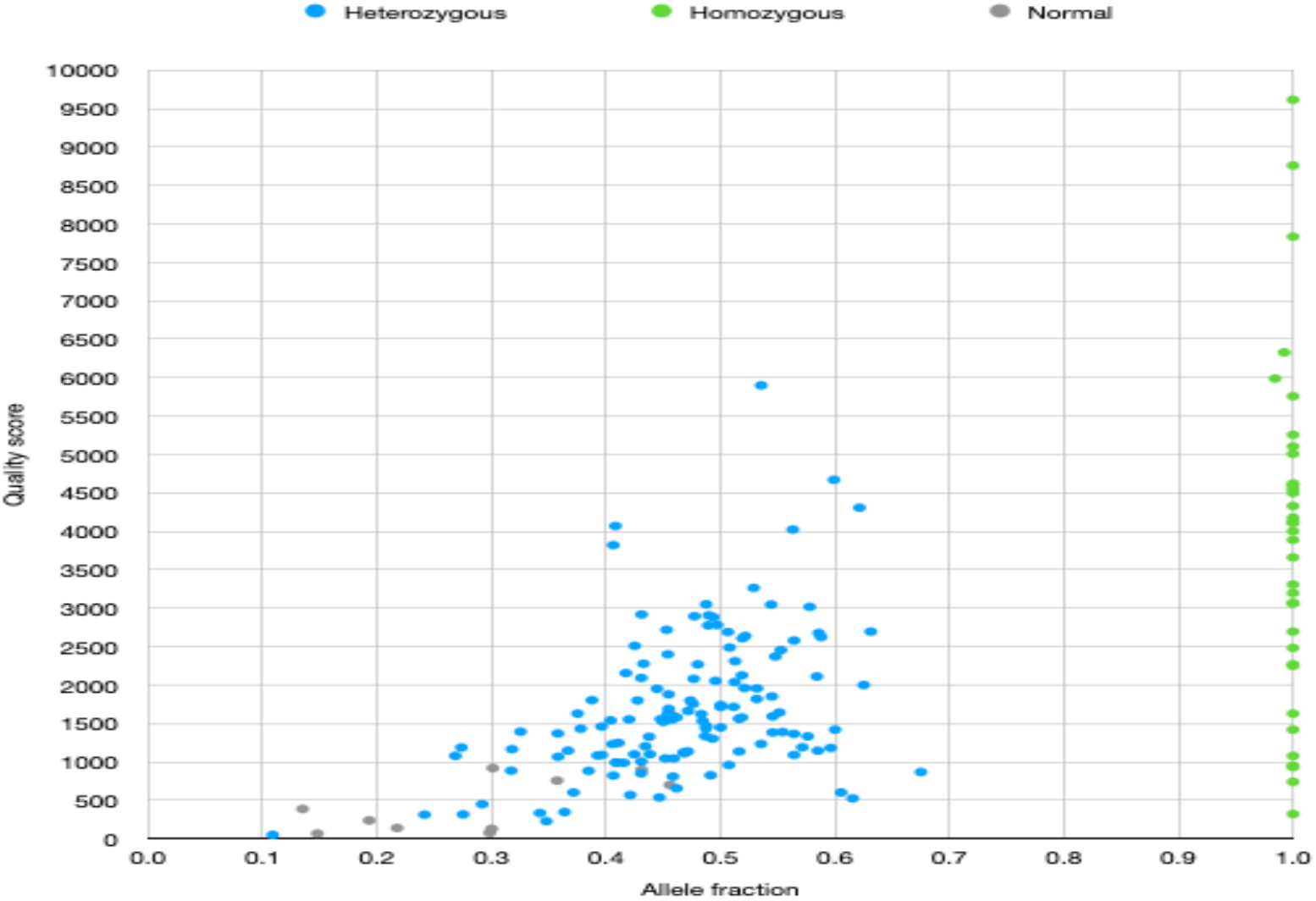
Distribution of indels according to the quality score and allele fraction

## Discussion

With the ability to simultaneously sequence more than 5000 disease-associated genes in a quick and cost-effective fashion, NGS has been supplanted Sanger sequencing as the method of choice in the diagnostic field at the laboratory level (1). The high sensitivity and specificity of NGS results are inevitable as clinicians rely on the data to make clinical decisions including treatment, early intervention, surveillance, and follow-up examination. Reporting a false-positive variant can have a significant impact on the health care management for patients and their families. Therefore, secondary validation, such as Sanger sequencing, is necessary to confirm the reported variants. The College of American Pathologists (CAP) recommends performing confirmatory testing for identified variants from the NGS, commonly using Sanger sequencing (2). However, the cost and time required to validate variants detected in NGS data using Sanger sequencing can hinder a timely diagnosis. Technical confirmation of high-throughput NGS data from relatively low-throughput sequencing prior to reporting is rarely done for other types of molecular testing. Furthermore, CAP suggests that the laboratory can have the flexibility to not perform Sanger sequencing if it has the criteria to support the high-quality NGS results that did not require Sanger sequencing (5). In the present study, we showed that all SNVs at >250 quality scores and 0.3> allele fractions were confirmed by Sanger sequencing. Although indels might also have a quality threshold, there are concerns about not enforcing Sanger sequencing due to the small number of sample size, indels base pairs size, repeat sequence, raw data alignment. Thus, we suggest that exome-derived SNVs with >250 quality scores and 0.3> allele fractions are not routinely required for Sanger validation.

Multiple studies have demonstrated that the quality threshold based on NGS-derived variants confirmed by Sanger sequencing is determined in the clinical diagnostic setting as in the present study (3, 8–10, 12). Several laboratories, including those in academic and commercial areas, have conducted studies to determine the quality threshold according to the various types of target captured panels or exome, different sequencers, and sample sizes ranging from dozens to thousands, resulting in finding minimum quality thresholds that met nearly 100% Sanger confirmation depending on each laboratory (3, 8–10, 12). Another study showed that 5,798 SNVs from the 684 exome dataset that met the quality threshold for variant calling were validated by Sanger sequencing, of which even 17 variants were confirmed by Sanger sequencing using newly designed primers due to the failure of first Sanger sequencing (11). High quality variants located on homo-polymeric regions larger than seven bases, homopolymer at the 100bp flanking sequencing, or having pseudogenes were not detected by Sanger sequencing, but were identified by mass spectrometry mapping (8). The findings from our study were almost concordant with those of previous studies. Additionally, four variants, including three SNVs and one indels, were not confirmed bi-directionally by Sanger sequencing due to reverse poly A or poly T sequence in the present study. These variants were clearly read on the raw .bam file and identified on the forward Sanger sequencing, and eventually reported to the clinicians. Collectively, these results can suggest that routine Sanger confirmation for NGS-derived variants is not essential due to the high accuracy of variant calling process and the possibility that incorrectly reading true positive variants.

The inclusion of Sanger confirmation of WES in the clinical diagnostic setting requires at least one more week due to the additional steps to the experiment process and subsequent costs, which are generally estimated to be US $240 per one variant by other laboratories and US $99 per one variant by our group (3). When the first Sanger sequencing fails, additional processes, including designing new primers, ordering, and re-running should be performed, causing a delay in reporting to clinicians and patients. Approximately 70% of identified variants from WES did not need to perform Sanger confirmation according to our minimum quality threshold and also had a reduced turnaround time, which can lead to reporting to clinicians within 3-4 weeks.

However, Sanger sequencing for indels may be necessary to define the exact genomic location or confirm quality assurance. In many cases, a single indel is listed as multiple different variant calls by NGS, of which the allele fraction or read depth is low (9). In this study, most indels proven to be false positive had low allele fraction and were one of multiple different variant calls despite a high quality score. Four variants (two unique variant) were not identified in the raw .bam file. Therefore, the raw .bam file should be reviewed in regions where indels or multiple calls for indels occur, and then, Sanger sequencing should be performed for accurate variant identification.

In conclusion, our results indicated that Sanger confirmation is not necessary for exome-derived SNVs with >250 quality scores and 0.3> allele fractions set to an appropriate quality threshold. All indels and SNVs that do not meet the quality threshold should be reviewed by raw .bam file and Sanger confirmation should be performed to ensure accurate reporting. Furthermore, the current quality threshold should be updated to improve quality assurance and develop stringent standards, taking into account that this threshold was set for a small sample size and simple features.

